# Ectopic expression of two cone opsins in mouse RGCs results in opposite responses to light stimulation, likely due to differential G protein activation

**DOI:** 10.1101/2025.09.10.675309

**Authors:** Marco Rucli, Nika Vrabic, Corentin Joffrois, Antoine Chaffiol, Melissa Desrosiers, Serge Picaud, Olivier Marre, Stefan Herlitze, Deniz Dalkara

## Abstract

Retinitis pigmentosa, a leading cause of inherited blindness, results in photoreceptor degeneration that current optogenetic approaches aim to address through microbial opsin expression in retinal ganglion cells (RGCs). While these microbial proteins restore light sensitivity, their clinical potential remains limited by low light sensitivity and immunogenicity risks. Recent efforts have focused on vertebrate opsins as safer alternatives, with mid-wavelength cone opsin (MW-opsin) demonstrating RGC depolarization via endogenous G-protein signaling. In this study, we reveal a surprising divergence in signaling outcomes of expressing short-wavelength mouse cone opsin (Opn1sw) in RGCs of blind mice. Using multielectrode array recordings and whole-cell patch clamping, we demonstrate that Opn1sw induces membrane hyperpolarization in RGCs – a stark contrast to MW-opsin’s depolarizing effects. This unexpected inversion suggests differential engagement of intracellular signaling pathways, potentially stemming from distinct G-protein coupling preferences. Comparative analysis of native G-protein expression profiles in RGCs versus cone photoreceptors supports this hypothesis, revealing mismatches that may explain ectopic opsin behavior. Our findings challenge the assumption of conserved opsin signaling across spectral subtypes and cell types, highlighting critical gaps in understanding vertebrate opsin-G protein interactions in non-native cellular environments. This discovery points to the necessity of systematic characterization of opsin signaling networks in target retinal cells, a prerequisite for engineering optimized optogenetic tools that reliably produce desired electrophysiological outcomes. By establishing spectral subtype-dependent signaling divergence, our work redefines parameters for developing next-generation vision restoration therapies.

## Introduction

Inherited retinal diseases affect about 2 million people worldwide and are the main cause of blindness in the adult population of industrialized countries(*1*). The main group of such diseases is retinitis pigmentosa (RP), a progressive degenerative disease affecting photoreceptors and causing retinal degeneration(*2*). At late stages of degeneration, when photoreceptors are completely lost, the best opportunity to restore light perception is by optogenetic strategies, which transform surviving inner retinal neurons into light-sensitive cells. Over the past 20 years, diverse attempts have been made at achieving this goal(*3*). Microbial opsins, light-sensitive proteins found in algae and bacteria(*4*) demonstrated their ability to efficiently reactivate retinal ganglion cells (RGCs), bipolar cells (BCs) and AII amacrine cells (AII ACs) upon light stimulation with great temporal resolution(*5–8*). Microbial opsins function as light-gated ion channels or pumps that directly transform light into electrical function as light-gated ion channels or pumps that directly transform light into electrical current without the need for G protein signaling. This process begins with light absorption by a chromophore, typically an all-trans-retinal form in the dark state, which is covalently bound to the opsin protein. When a photon is absorbed, the chromophore isomerizes to a cis-retinal form, inducing a conformational change in the opsin(*9, 10*). This structural shift leads to the opening of an ion channel or the activation of an ion pump, depending on the specific type of microbial opsin. Once activated, these channels or pumps allow the flow of specific ions (such as protons, sodium, or chloride) across the cell membrane, generating an electrical current. This process occurs with millisecond-scale response times, allowing for precise temporal control of ion flow. After activation, the opsin eventually returns to its original state, either through thermal relaxation or, in some cases, through exposure to a different wavelength of light. The direct light-to-current conversion mechanism of microbial opsins makes them particularly useful for optogenetic applications, enabling rapid and precise control of neuronal activity in response to light stimulation. These opsins, however, generally lack the sensitivity and the adaptability of vertebrate opsins and require high light intensities to be activated (which can translate to patients needing to wear light-amplifying goggles(*11*)). To improve on this aspect, researchers turned to G-protein coupled opsins, such as rhodopsin, melanopsin and cone opsins (expressed in rods, in intrinsically photosensitive retinal ganglion cells (ipRGCs) and in cones, respectively). These opsins are characterized by greater light sensitivity due to their reliance on a signal-amplifying G protein cascade(*12*). Melanopsin and cone opsin signaling differ in several key aspects. Melanopsin, more closely related to invertebrate visual opsins, couples to G(q)-type proteins and activates phospholipase C, leading to the opening of TRP-channels. This signaling cascade results in more sustained responses with post-stimulus persistence, contrasting with the transient responses of cone opsins. Melanopsin requires higher light intensities for activation and primarily functions in non-image-forming processes like circadian photoentrainment and pupillary light reflex. It is expressed in intrinsically photosensitive RGCs of the inner retina. Conversely, cone opsins, found in cone photoreceptors of the outer retina, are involved in visual perception and color vision, with lower activation thresholds. They typically activate a different G-protein cascade and exhibit faster constriction latencies. Unlike cone opsins, melanopsin retains its chromophore following photoactivation, similar to invertebrate opsins. These distinctions in signaling mechanisms and characteristics enable melanopsin and cone opsins to contribute uniquely to various aspects of light detection and visual processing in vertebrates and have been useful in their implementation as optogenetic tools to study neural circuits(*15*). Moreover, ectopic expression of vertebrate opsins in inner retinal neurons generally resulted in highly sensitive light perception restoration, but both melanopsin and rhodopsin generated responses that are considered too slow for successful patterned vision restoration(*16–18*). The medium-wavelength sensitive opsin (MW-opsin) from cones, on the other hand, proved both fast and sensitive when ectopically expressed in mouse RGCs, actually combining the advantages of microbial opsins and vertebrate opsins and resulting in effective vision restoration in blind mice(*17*). Using human MW opsin in mouse RGCs, Berry and coworkers observed robust depolarization and increased firing rates in response to light. This effect was surprising because MW opsin, like other cone opsins, is expected to couple to Gi/o signaling pathways, which typically lead to hyperpolarization. The depolarizing outcome likely arises from unique aspects of the intracellular environment in RGCs: the specific repertoire of available G protein subunits, downstream effectors, or ionic homeostasis may allow MW opsin to activate alternative G protein pathways (such as Gq/11), or it may cause secondary signaling effects that ultimately increase excitability—contrasting sharply with the canonical inhibitory Gi/o signaling seen in other contexts. Thus, despite using the same fundamental signaling mechanism as other opsins, MW opsin in RGCs produced an excitatory, rather than inhibitory, light response in these prior studies. While thoroughly describing MW-opsin’s properties, the study did not investigate the molecular mechanisms responsible for such behavior. It is known, however, that GPCRs are selective towards specific G protein subtypes(*19*), and that different cell types express different subunits of such proteins(*20*). Investigating the molecular details of how these opsins work as optogenetic tools is needed to better understand and use them to restore vision. To do so, we decided to test a second vertebrate cone opsin, the mouse short wavelength-sensitive opsin Opn1sw, in mice RGCs, as it was previously reported to efficiently activate GIRK channels through the G_i/o_ pathway both *in vitro* and *in vivo*(*21*).

We found that Opn1sw triggers RGC hyperpolarization upon light stimulation with both UV and visible light, a contrasting result to the previously reported vertebrate-opsin mediated responses in RGCs which were all depolarizing(*16, 22*). We show that the G protein subunits, which are a fundamental element of the opsin cascade, greatly differ between RGCs and cone photoreceptors. We speculate that the opposite effects induced by the medium-wavelength sensitive opsin and Opn1sw on membrane polarity are the result of the activation of different G protein cascades in RGCs. Additionally, by comparing Opn1sw to the microbial type hyperpolarizing opsin GtACR1, we show that both opsins offer similar light sensitivity but the former is considerably deficient in terms of temporal resolution. Finally, our behavioral tests show that SNCG-Opn1sw is not capable of restoring light perception in blind animals under dim light conditions, potentially due to the short wavelength light reaching the retina less efficiently through the mouse eye’s lens in vivo. Overall, our results hint towards the need for a deeper understanding of the intracellular pathways activated by ectopically expressed vertebrate opsins within non-photoreceptor cells of the retina. This is a *sine que non* condition for designing opsins for vision restoration. Moreover, the improvements in the sensitivity of microbial opsins such as GtACR1 currently allow them to operate at similar light sensitivity with greater time resolution and predictability making microbial opsins the preferred opsins for visual restoration attempts.

## Materials and methods

### Animals

For in vivo and ex vivo experiments, C57BL/6J (WT) and C3H/FeJ (alternative names: rd1nobfree or rd1) mice were used. C57BL/6J mice were purchased from Janvier Labs, while C3H/FeJ mice were initially purchased from Jackson Laboratory and maintained as a colony at the Vision Institute. C3H/FeJ mice are characterized by a homozygous mutation in the Pde6b gene, causing rapid retinal degeneration and complete photoreceptor loss by approx. PND90(*23*). In addition, C3H/FeJ mice used in this project did not carry the mutated Gpr179 gene giving the nob (“no b-wave”) phenotype, ensuring that any residual activity from photoreceptors would be normally transmitted across the retina(*24*).

### Plasmid design and AAV production

The Opn1sw-tdTomato transgene was amplified from existing internal plasmids and manually cloned into am SNCG-containing plasmid via the In-Fusion cloning method. Since the backbone DNA located between the promoter and the WPRE sequence was flanked by restriction enzyme sites, the plasmid was incubated with restriction enzymes in order to remove the sequence to be replaced. The transgene was then PCR-amplified and incubated with the open backbone and a recombinase, allowing for the final plasmid to be synthesized. The final plasmid structure was organized as following: ITR-SNCG-SV40-Opn1sw-tdTomato-WPRE-BGHpA-ITR. The plasmids carrying ITR-SNCG-SV40-mMWO-GFP-WPRE-BGHpA-ITR and ITR-SNCG-SV40-GtACR1-mCherry-WPRE-BGHpA-ITR were purchased from Vectorbuilder (https://en.vectorbuilder.com/). The final plasmids were used to produce the desired AAV vectors in-house using the plasmid co-transfection method: vectors were purified with iodixanol ultracentrifugation, and titer was obtained by real-time PCR using the standard curve method for absolute quantification. Final titers were 4.98e+14 vg/ml, 3.57e+14, 3.91e+12 vg/ml for SNCG-Opn1sw, SNCG-Opn1mw and SNCG-GtACR1, respectively.

### AAV injection and fundus imaging

All animal studies were conducted in accordance with protocols approved by an ethical committee (submission n°30129 and n°40310) and following the ARVO Statement for the Use of Animals in Ophthalmic and Vision Research. Animals were injected intravitreally between PND30 and PND60 with 2µL of viral solution. Prior to injection, eyes were dilated with Mydriaticum 0.5% (Théa) and Néosynéphrine 10% (Europhta). Eyes were then coveredwith Lubrithal (VetXX), a protective gel. Mice were anesthetized via Isofluorin^®^ inhalation (5% induction and 2% sustain). Fundus images were taken either two weeks (subretinal injection) or four weeks (intravitreal injection) post injection using a fundus camera (Micron IV; Phoenix Research Lab) equipped with red and green filters for fluorescence detection.

### Immunihistochemistry

To obtain retinal sections, mice were euthanized with CO_2_, followed by cervical dislocation. Eyes were collected, put in a 4% paraformaldehyde-PBS solution for 2h at room temperature and then washed three times with PBS 1X. Eyes were then dissected, a procedure needed to eliminate the lens and the cornea, leaving just the eyecup. Eyecups were put in a 10% sucrose-PBS solution for 1h and then in a 30% sucrose-PBS solution overnight. The following day, eyecups were washed in liquid tissue freezing medium (Microm Microtech) at room temperature and frozen in the same medium using liquid nitrogen. 12 μm thick slices were cut using a cryostat (Leica CM3050S), they were placed on glass slides (Thermo Scientific) and then stored at −20°C. For immunostaining, prior to incubation with the blocking solution, slides were unfrozen for 30 minutes and then washed three times for 5 minutes with PBS 1X. Slides were then incubated for 1h at RT in a blocking buffer (PBS-BSA 1%-Tween 0.1%-Triton X-1000.1%) with and DAPI (1:2000); no antibodies were used for fluorescent protein detection. Lastly, slides were again washed three times for 5 minutes in PBS 1X, mounted in Fluoromount Vectashield (Vector Laboratories) and covered with a glass coverslip. Slides were imaged via laser-confocal microscopy (Olympus IX81) and the analyzed with FIJI software.

### Single cell RNA sequencing

#### Preparation of single-cell suspensions from mouse retinas

Following cervical dislocation, eyes from WT and rd1 mice aged PND120 to PND190 were enucleated and transferred to CO2-independent medium (Thermo Fisher Scientific, Ref. 18045088) for dissection. The retinas were isolated, chopped into small fragments, and washed twice in warm homemade Ringer solution without Ca2+. The fragments were then incubated in 1 ml of activated papain (4 units/ml, Worthington Biochemical Corp., Ref. LSO 3124) at 37°C for 20 minutes. Digestion was halted by adding 1 ml of Neurobasal-A medium supplemented with L-Glutamine (1%) and fetal bovine serum (5%). DNase I (0.02 mg/ml, Sigma-Aldrich, Ref. D4263) was then added, and the tissue was gently dissociated by trituration using a 1000 μl pipette tips. A brief centrifugation (30 seconds, 115 g) was performed to separate the dissociated cells (supernatant) from aggregates. This process was repeated 4 to 6 times until the retinas were fully dissociated. The total cell suspension was then centrifuged for 10 minutes at 115 g. The resulting pellet was resuspended in a small volume of 1X PBS (calcium and magnesium-free) containing non-acetylated bovine serum albumin (400 μg/ml), then viability and cell concentration was determined using a Malassez counting chamber.

#### Single-cell RNA sequencing and Bioinformatic analysis

Single cells were sorted using the Chromium X controller (10 x Genomics) and used to construct scRNA libraries in accordance with Chromium Next GEM Single Cell 3′ Library v3.1 and dual Index Kit TT. The Illumina Novaseq 6000 sequencer was used to sequence the libraries. The fastq obtained after sequencing were preprocessed using the cellranger pipeline (version 7.0.1, reference genome: cellranger reference dataset refdata-gex-mm10-2020-A) with default options; the filtered count matrices were then used for the analyses. All datasets were integrated using the R package Seurat (*25*), following their vignettes. After the integration, the standard pipeline was used to filter (percent.mt < 10, nFeature_RNA > 200, nCount_RNA > 1000), cluster (resolution = 1) and create the UMAP/tSNE visualizations using the default values. The resulting dataset was annotated with other datasets: one from(*26*), the RGC dataset from(*27*), the amacrine cells dataset from (*28*) using the cell type classification method from the Seurat package. The final dataset was then converted to an anndata file for visualization in cellxgene(*29*) with the cellxgene_vip plugin(*30*) for further analyses.

### Electrophysiology

MEA recordings were performed on WT and rd1 mice at >PND90 and at least 4 weeks post injection. Mice were euthanized with CO2, followed by cervical dislocation. Eyes were collected and rapidly dissected under dim red light in oxygenated Ames media. Retinas were then fixed to a supporting cellulose membrane, previously treated with 0.1% (w/v) in H2O Poly-L-lysine solution (Sigma) for 1h, and gently placed, ganglion cell facing down, in the recording chamber of a 256-channel MEA system (MEA256 - electrode size 30 microns and spacing between electrodes 100 microns; Multichannel Systems). The retina was provided with constant perfusion of oxygenated Ames media at 34°C at a rate of 1-2mL/min, with the addition of 20μM 9-cis retinal (Merck). The metabotropic glutamate receptor agonist L-AP4 (Tocris Bioscience) was added to the fresh medium when testing HKamac-Opn1sw-injected animals at a concentration of 50μM.

Full field light stimuli were generated using a Polychrome V monochromator (Olympus)controlled by the STG2008 generator (MCS) at intensities ranging from 10^13^ to 10^16^ photons*s-1*cm-2. When only one wavelength was used, the experimental protocol consisted in 5 repetitions of 1s stimuli with 60s dark intervals, unless otherwise specified. For the spectral analyses, we used a set of wavelengths ranging from 300nm to 600nm in steps of 50nm, to which the melanopsin’s peak absorption wavelength (480 nm) was added; furthermore, the peak absorption wavelength for Opn1sw added (380 nm). In the spectral analysis, the experimental protocol consisted in 1s stimuli with 30s dark intervals, with 5 repetitions for each wavelength.

### MEA data analysis

Raw RGC electrical activity was amplified, sampled at 20 kHz with a 200 Hz high-pass filter and recorded using MC_rack software (Multi Channel Systems) for further offline analysis. Firing rate was calculated using 10ms bins over 5 repetitions, unless otherwise specified. We then estimated the mean firing rate and the SD prior to the stimulus. For fast depolarizing cells, we considered that a response was generated if it exceeded the mean firing rate prior to the flash by at least 3 times the value of SD during the light stimulation. Similarly, we considered that a hyperpolarizing response was generated if it was lower than the mean by at least 3 times the SD in the 2s following the light stimulation. The late detection window was motivated by the fact that SNCG-Opn1sw responses peaked after the stimulus offset. To calculate the percentage of responding electrodes, we divided the number of responding electrodes by the number of active electrodes (we considered that an electrode was active if it recorded at least 100 spikes over the course of the recording). For slowly depolarizing cells, we considered that a response was generated if it exceeded the mean by at least 3 times the value of SD in the 10s following light stimulation. For intensity-response analyses, mean values for each intensity were normalized to the highest response.

### Light-avoidance behavior

We tested in vivo light avoidance behavior with a light/dark box (Panlab) consisting of two equal-sized compartments separated by a non-transparent wall with a 7 cm x 5 cm hole in the middle. Illumination was provided through white or wavelength-specific LED PCBs (Thorlabs) which were singularly soldered to LED connection cables and powered through a T-cube TM LED driver (Thorlabs). LEDs were placed in the center of the top part of the light compartment; the light intensity was measured approximately at the level of the mice’s eyes in the center of the light compartment through a spectrophotometer (Thorlabs). The day prior to testing, animals were let acclimate for 45min in the box at ambient lighting conditions, then dark-adapted overnight and tested on the following day. During the test, animals were put in the illuminated compartment and given 3 min to discover the dark compartment: a 15min test started as soon as the mouse completely entered the dark compartment. Mice failing to reach the dark compartment within 3 min were excluded from the analysis. The two compartments were intensively cleaned before each mouse began the test. Animal movements were recorded with a camera and the time spent in each compartment was manually computed, using the center of the mice’s body to determine which compartment it occupied. Statistical differences between groups were assessed via one-way ANOVA test (statistical significance set at P<0.05).

## Results

### AAV injection leads to strong and cell type specific expression of Opn1sw and Opn1mw in mouse RGCs

Previous studies showed that vertebrate opsins can be successfully implemented as optogenetic tools for vision restoration, although only the medium wavelength-sensitive opsin proved fast enough to allow for patterned vision restoration(*16, 17*). Given the ability of the medium wavelength-sensitive opsin to trigger rapid depolarizing responses(*22*), we asked what type of response would be mediated by the other mouse cone opsin, the short wavelength-sensitive opsin Opn1sw, when expressed in RGCs. We therefore tested Opn1mw alongside Opn1sw in order to have a direct comparison between Opn1sw and the benchmark depolarizing vertebrate opsin. Opn1sw-tdTomato and Opn1mw-GFP were cloned downstream of the strong and RGC-specific promoter SNCG(*31*) (**Figure 1A**), packaged into AAV2 capsids and injected intravitreally at equal doses into the eyes of rd1 mice, displaying fast retinal degeneration(*32*). Four weeks after injection, the fundus images showed pan-retinal expression for both opsins, although brighter expression was noted for Opn1sw tagged with tdTomato (**Figure 1 B and D**). Both opsins were expressed exclusively in RGCs based on cross sections of the retina (**Figure 1C and E**): Opn1mw showed partial cytoplasmic expression, while Opn1sw showed stronger membrane-bound and axonal expression. After ensuring expression in RGCs for both opsins, we further investigated their behavior using electrophysiology.

**Figure 1.**
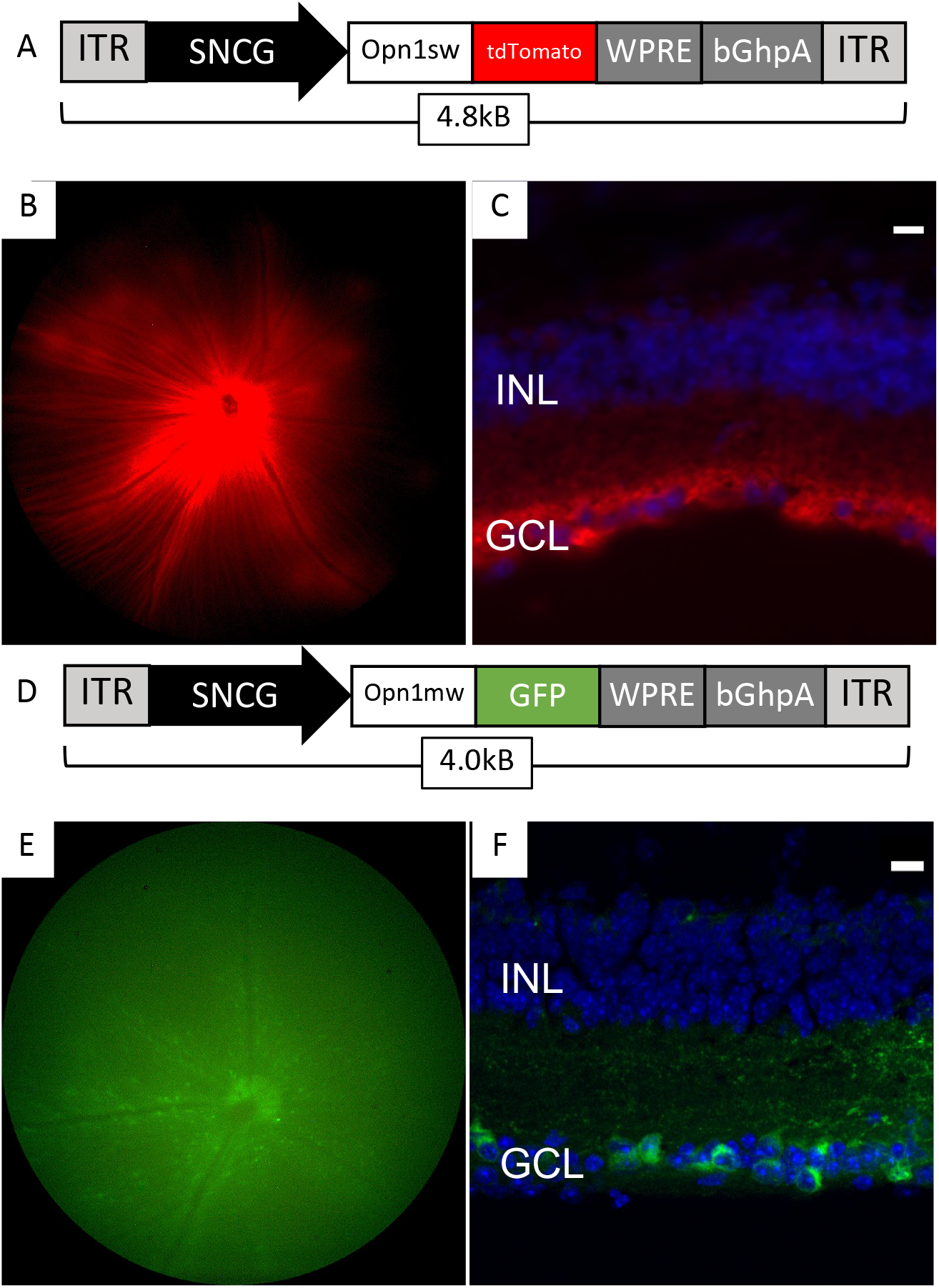
Opn1sw and Opn1mw expression in rd1 RGCs. A) Schematic representation of AAV expression cassette for SNCG-Opn1sw. B) Transgene expression in fundus images 4 weeks post intravitreal injection C) Cross sections from the injected retina 4 weeks post injection showing cellular localisation of Opn1sw in red. D) Schematic representation of AAV expression cassette for SNCG-Opn1mw. E) Transgene expression in fundus images 4 weeks post intravitreal injection F) Cross sections from the injected rd1 retina 4 weeks post injection showing cellular localisation of Opn1sw in green. : 10µm.

### Opn1sw and Opn1mw mediate opposite responses in RGCs of the rd1 retina

To evaluate opsin-mediated responses in the degenerated rd1 retina, we performed multielectrode array (MEA) tests on Opn1sw- and Opn1mw-injected animals. As expected, SNCG-Opn1mw-injected retinas displayed RGC depolarization when illuminated with 535nm green light (**Figure 2A**)), as previously reported(*22*). We observed, however, a poor efficacy of SNCG-Opn1mw under our experimental conditions as a maximum of 6% of the active MEA electrodes(that is, electrodes displaying spiking activity during the experiment) displayed such behavior (achieved at ~1e+14 photons*cm-2*s-1) (**Figure 2B**). We attribute such low value to the partial mis-localization of the Opn1mw protein in the cytoplasm (**Figure 1C**). Another reason for such low efficacy could lie in the notably poor contact between MEA and RGCs around optic nerve, which did not always allow for the recording from the highest transgene-expressing region of the retina in the case of Opn1mw.

**Figure 2.**
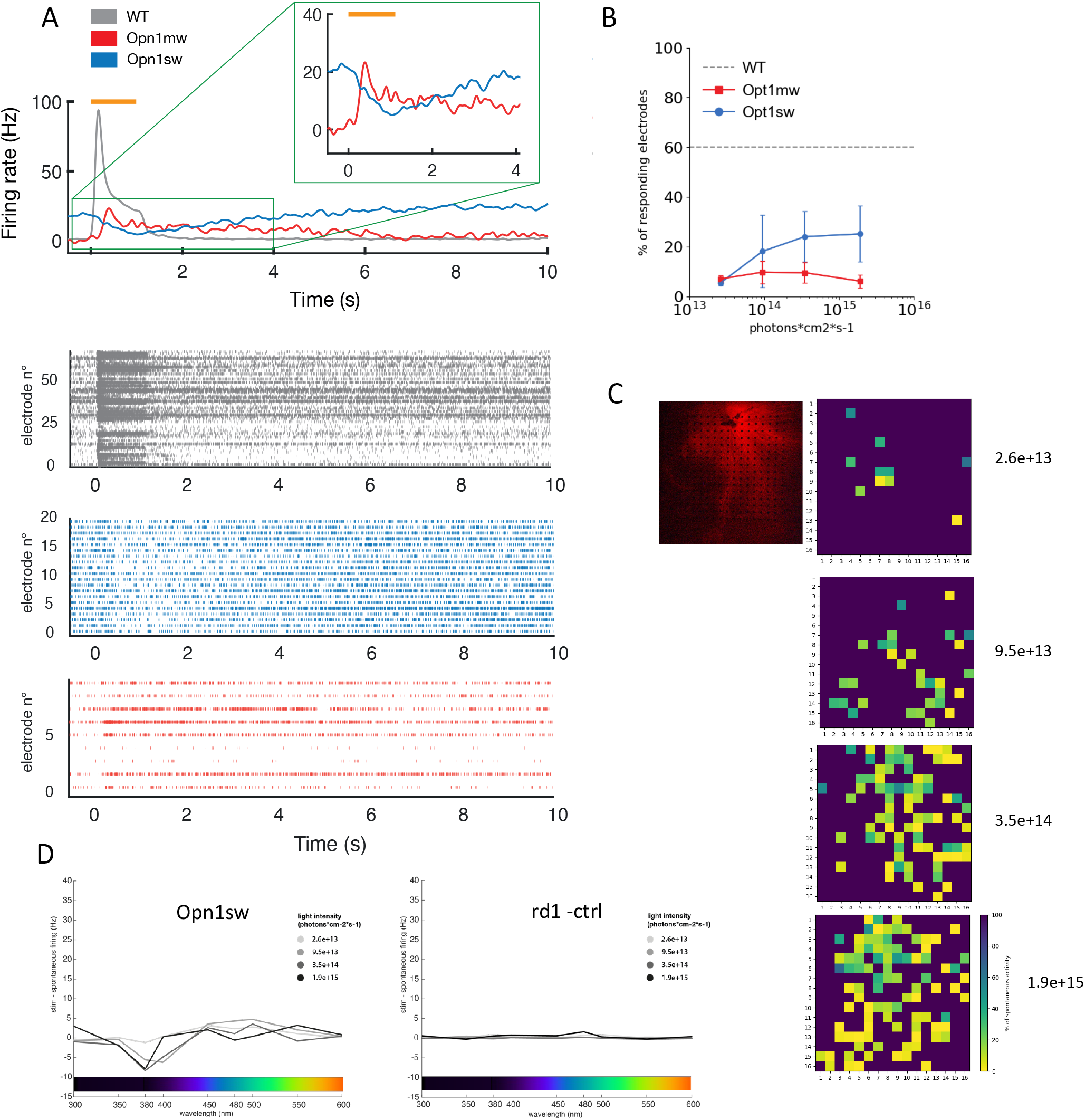
Opn1sw- and Opn1mw-mediated responses in RGCs. A) (Top) Average RGC response of WT (grey), Opn1sw-injected (blue) and Opn1mw-injected (red) rd1 retinas. (Bottom) Raster plot of average RGC responses to 5 flashes of 1s at 380 nm light (WT, 64 electrodes; Opn1sw, 20 electrodes) and 535 nm light (Opn1mw, 9 electrodes). Light intensity: 3.5e+14 photons*s-1*cm-2. B) Proportion of responding electrodes across different light intensities in Opn1sw-injected (blue dots, N = 3) and Opn1mw-injected retinas (red squares, N = 2). C)Fluorescence image of an injected retina overlayed on the MEA (left) Representative heatmaps of Opn1sw-mediated hyperpolarizing responses in the RGS of the rd1 retina shown on the left panel across multiple light intensities. Mean values across 5 flashes of 1s at 380 nm were averaged to the strongest hyperpolarizing response (right). D) Representative spectral analysis at multiple light intensities in Opn1sw-injected and in uninjected rd1 retinas with a peak hyperpolarizing response at 380 nm in Opn1sw-injected retinas (N = 43, 19, 41, 4 electrodes at 2.6e+13, 9.5e+13, 3.5e+14, 1.9e+15 photons*s-1*cm-2, respectively) but not in uninjected controls (N = 31, 167, 93, 39 electrodes at 2.6e+13, 9.5e+13, 3.5e+14, 1.9e+15 photons*s-1*cm-2, respectively); retinas were presented with flashes between 300 nm and 600 nm in steps of 50 nm, with the addition of 380 nm and 480 nm (corresponding to the peak absorption of Opn1sw and melanopsin, respectively); 5 flashes were displayed at each wavelength.

Contrary to our expectations, when Opn1sw-injected retinas were illuminated with a full-field flash at 380nm light (close to the opsin’s peak absorption), we observed a strong cellular hyperpolarization (**Figure 2A and C**). The time to peak (time to reach the maximal hyperpolarization) was quantified to be 1.12 ± 0.37 s (n=9 retinas). Opn1sw-induced hyperpolarizing responses were detectable at light intensities as low as ~2,3e+13 photons*cm^−2^*s^−1^, and the proportion of hyperpolarizing electrodes increased at increasing light intensities, reaching a plateau at ~3,5e+14 photons*cm-2*s-1 (**Figure 2B**); at this intensity, about 25% of electrodes showed hyperpolarizing responses, while in wild-type retinas the depolarizing events following photoreceptor activation were observed in about 60% of electrodes (**Figure 2B**, dashed line). Furthermore, we confirmed the presence of Opn1sw-mediated RGC hyperpolarizations at a single cell level through spike sorting (**Supplementary Figure 1**). In order to confirm that the observed responses were generated by Opn1sw, we performed a spectral analysis in both SNCG-Opn1sw-injected and uninjected rd1 control retinas. As expected, spectral analysis showed a peak hyperpolarization at 380nm in Opn1sw-injected retinal but not in uninjected controls, with a reduced but still detectable activation at 400nm (**Figure 2D**), in agreement with Opn1sw’s spectrum and with previously published data in HEK293 cells^21^. Once again, hyperpolarization became stronger at increasing light intensities (**Figure 2D**). Overall, Opn1sw-induced response came as a surprising result, as all previously published data on vertebrate opsins described an opposite, depolarizing effect being generated by the activation of the opsin in RGCs^15,16^. These results hint towards a selective activation of different intracellular G protein-coupled pathways by Opn1sw and Opn1mw, likely determined by the differences in their intracellular domains (ILs and C-terminal).

In parallel to the diversification of the vertebrate opsin toolbox for control of neuronal activity in different contexts^26–29^,there has been significant developments in engineering microbial opsins for better light sensitivity^30^. We thus decided to test Opn1sw against the light-gated hyperpolarizing chloride channel GtACR1^31^, which previously showed to efficiently hyperpolarize HEK293 cells^31^. When injected in rd1 mice, SNCG-GtACR1 showed good levels of expression 4 weeks post-intravitreal injection (**Figure 3A**). As expected, illumination with 515nm green light generated an extremely fast cellular inhibition of RGC’s spontaneous spiking, which was followed by a rebound depolarization (**Figure 3B**). Interestingly, GtACR1-induced hyperpolarization could be observed at intensities as low as 2.6e+13 photons*s-1*cm-2, which became stronger as light intensity increased, although the overall number of hyperpolarizing electrodes decreased (**Figure 3C**). The wider spread of hyperpolarizing electrodes at low intensities highlights once more the great ability of microbial opsins in generating a response independently of the cell (sub)type. Overall, SNCG-GtACR1 proved both faster and even more light-sensitive than SNCG-Opn1sw, indicating how wildtype vertebrate opsins are in many cases still inferior optogenetic tools for fast neuronal modulation when compared to microbial opsins.

**Figure 3.**
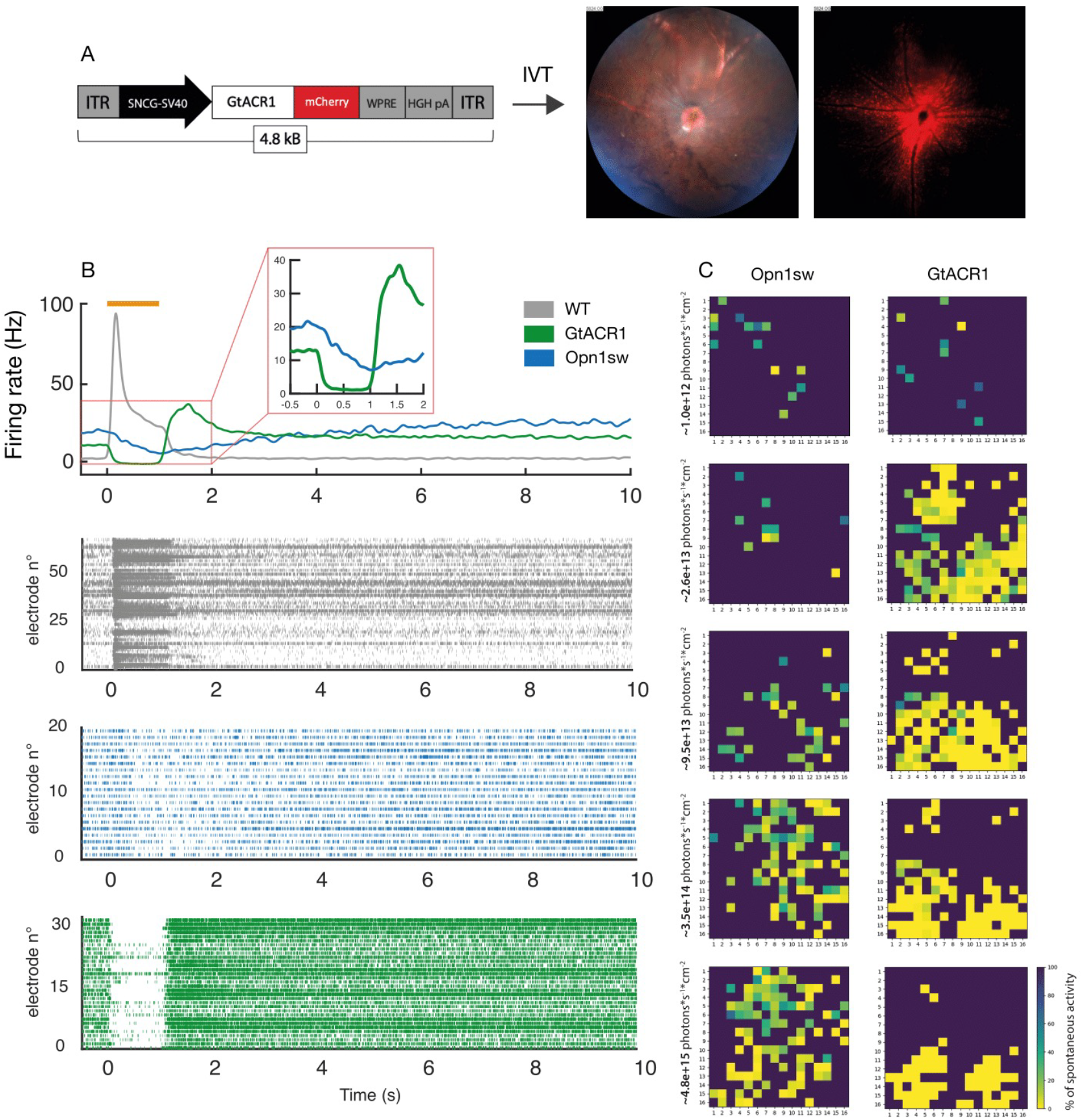
SNCG-GtACR1-mediated responses in RGCs. A) (Left) SNCG-GtACR1 expression cassette; (Right) Fundus imaging shows strong transgene expression four weeks after intravitreal injection. B) (Top) Average RGC response of WT (grey), Opn1sw-injected (blue) and GtACR1-injected (green) rd1 retinas. (Bottom) Raster plot of average RGC responses to 5 flashes of 1s at 380 nm light (WT, 64 electrodes; Opn1sw, 20 electrodes) and 515 nm light (GtACR1, 32 electrodes). Light intensity: 3.5e+14 photons*s-1*cm-2. C) Representative heatmap of GtACR1- and Opn1sw-mediated hyperpolarizing responses in the RGS of an injected rd1 retina across multiple light intensities. Mean values across 5 flashes of 1s at 380 nm (Opn1sw) and 515 nm (GtaCR1) were normalized to the strongest hyperpolarizing response.

### scRNA sequencing reveals an RGC-specific G protein expression signature

Puzzled by the Opn1sw-mediated hyperpolarization of RGCs, and to further investigate the relationship between the observed cellular responses and the G protein profile of RGCs, we decided to perform single-cell RNA sequencing (sc RNAseq) in SNCG-Opn1sw-injected and SNCG-Opn1mw-injected rd1 retinas. First, sc RNAseq further confirmed our histological results, showing strong and specific targeting of RGCs, (**Figure 4A**). Given that there are at least 46 distinct types of RGCs in the mouse retina(*27*), we wanted to ensure that our pan-RGC promoter SNCG(*31*) is similarly active across all classes of RGCs. Indeed, except for subtypes represented by only a few cells, we found that both Opn1sw and Opn1mw were similarly expressed by almost all RGC subtypes (**Supplementary Figure 2**). This result indicates that any difference in the two opsins’ effect on membrane potential is likely not determined by a biased, RGC subtype-specific expression of the transgene, but rather by an opsin-specific G protein cascade activation. Indeed, when we performed sc RNAseq on whole WT retinas, we observed considerable differences in the G protein profile of RGCs compared to the one of cone photoreceptors (**Figure 4B**). In cones, as expected, the most expressed subunits were related to the phototransduction cascade: Gɑ_t-c_ (Gnat2 or cone transducin), Gβ_3_ (Gnb3) and G*γ*_14_ (Gngt2). This was not the case for RGCs, where the most highly expressed genes were Gɑ_s_ (Gnas), Gɑ_o1_ (Gnao1), Gɑ_q_ (Gnaq), Gβ_1_ (GNB3) and G*γ*_3_ (Gng3). In cones, we also observed the presence of the rod-specific genes Gɑ_t-c_ (Gnat1 or rod transducin) and G*γ*_1_ (Gngt1), a seemingly contradictory result which can be explained by rod OS mRNA contamination, a common issue in sc RNAseq in the retina^33^.Overall, sc RNA seq suggest the possibility that the RGC-specific G protein landscape can be differentially activated by Opn1sw and Opn1mw, leading to opposite effects on RGC response polarity.

**Figure 4.**
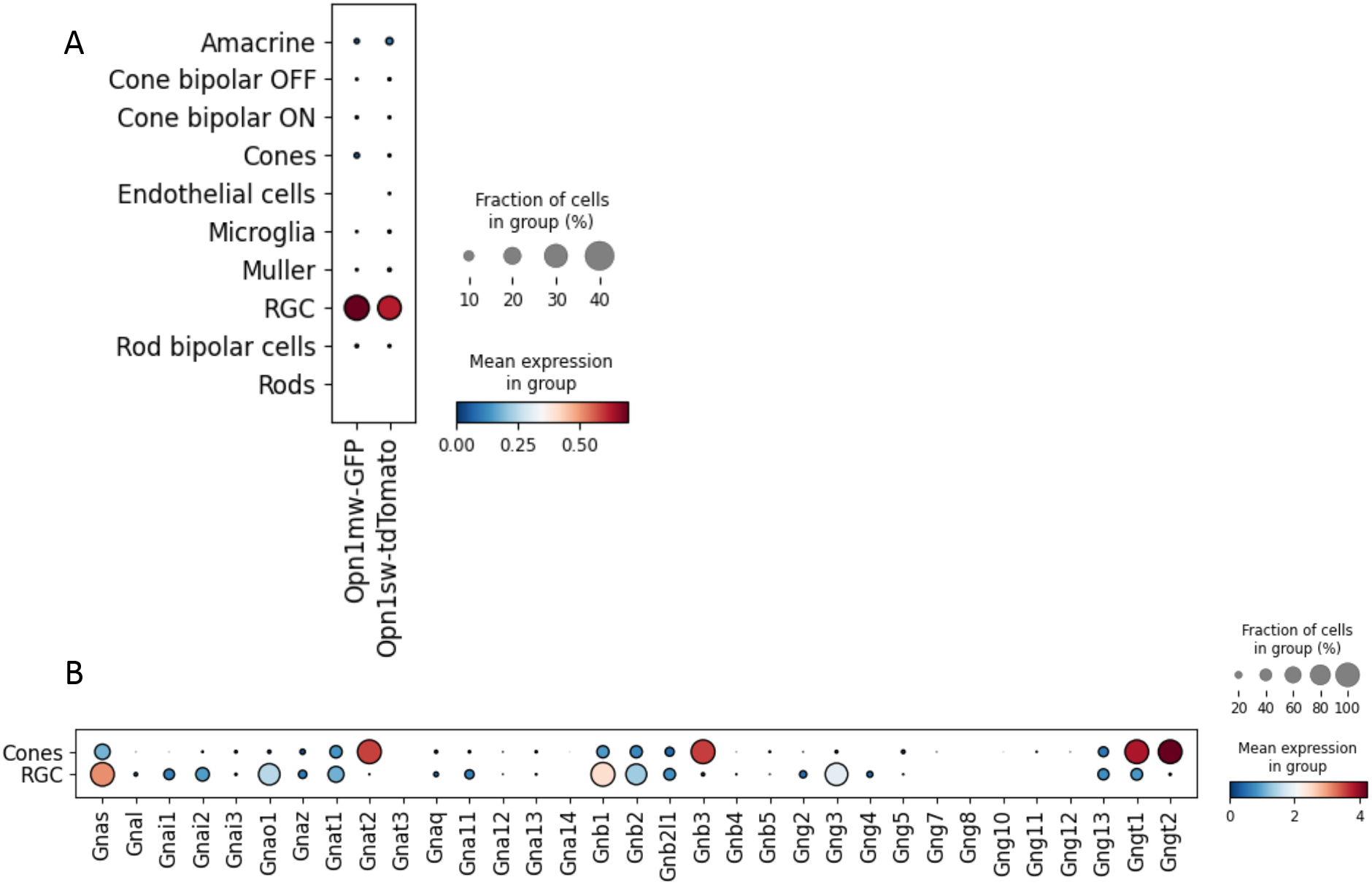
Single-cell RNA sequencing in WT and rd1 retinas. A) Sc RNA seq reveals good specificity of the SNCG promoter, which mainly targets RGCs when coupled to both Opn1sw and Opn1mw. B) The analysis of G protein-related genes in the WT retina reveals a major change in the expression between cone photoreceptors and retinal ganglion cells.

### Opn1sw is unable to restore light perception in vivo under low intensity white light

Encouraged by the behavioral responses previously reported for Opn1mw(*22*), we performed a simple the light/dark box test (**Figure 5A**) on animals expressing Opn1sw in their RGCs. This test was conducted to see if dim white light could elicit behavioral light avoidance responses *in vivo*, under the assumption that cellular hyperpolarization could nonetheless be interpreted by the visual cortex as a light-induced stimulus. When illuminated with white light at 100 μW*cm-2 or 10^14^ photons*s-1*cm-2 (the approximate illumination of an indoor office), unlike wild-type animals that spend 60-70% of their time in the dark compartment, SNCG-Opn1sw-injected animals showed similar behavior to uninjected negative controls, spending similar amounts of time in the two compartments (**Figure 5B**). Although the light intensity used in the behavioral test was comparable to the one that elicited responses in MEA experiments; this outcome is likely the result of the majority of the short wavelength light being filtered by the lens of the mouse eye in vivo. Indeed, the lens and cornea of the mouse retina are known to filter, especially blue light which is in the excitation maximum of Opn1sw and could thus be responsible for a lack of retinal response in our experimental conditions(*33*). Another potential explanation might be that the hyperpolarization of the blind retina is not able to induce a significant enough change in the light perception of the animal(*34, 35*). Alternatively, a strong interruption of the noisy background activity of retinal ganglion cells is potentially not processed as an event triggering light avoidance by the animal. Further testing with a broader range of intensities will be needed to elucidate this aspect of Opn1sw-induced hyperpolarization under in vivo conditions.

**Figure 5.**
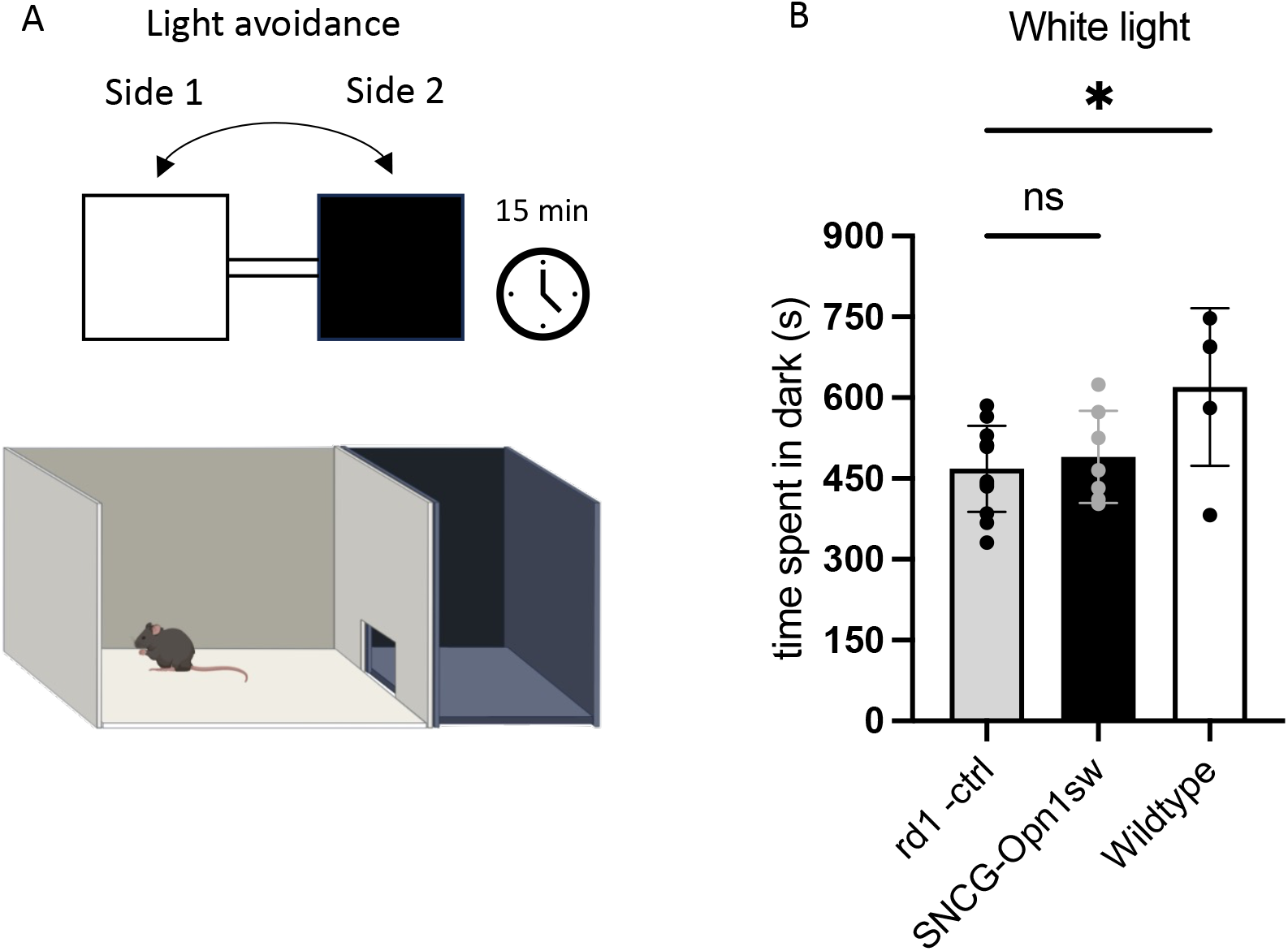
Behavioural analysis on SNCG-Opn1sw-injected rd1 animals. A) Schematic representation of the light-dark box experimental setup. B) Time spent in dark (light avoidance behaviour) for uninjected rd1 mice (grey), SNCG-Opn1sw-injected rd1 mice (black) and sighted wild-type mice (N = 12, 7, 5, respectively) at PND100. White light intensity: 100 μW cm-2 or approx. 2*10^14^photons*s-1*cm-2. Values are mean + SD. Statistical significance assessed using ordinary one-way Anova test: *adjusted *p* value < 0.05.

## Discussion

In recent years, vertebrate opsins have been thought to carry great potential as optogenetic tools for vision restoration purposes(*16, 18, 22, 36–38*). Compared to the first generation microbial-opsin based tools, vertebrate opsins are indeed more light sensitive(*21*); furthermore, MW-opsin-mediated responses successfully restored patterned vision in mice, indicating fast kinetics(*22*). Despite such undeniable progress, the exact molecular mechanism that is responsible for MW-opsin-induced responses remains elusive. This lack of knowledge is problematic for the development of novel therapies, as the activity of a vertebrate opsin intrinsically relies on downstream elements like G proteins and effector channels. To improve vertebrate opsins’ design for vision restoration purposes, it is therefore crucial to deepen our understanding of their interaction in ectopic contexts.

In this study, we tested a short wavelength sensitive mouse cone opsin, Opn1sw, against the already characterized Opn1mw. We found that Opn1sw induces sensitive RGC hyperpolarization upon light stimulation, an opposite effect compared to Opn1mw. Sc RNAseq revealed that transgene expression is homogenous across RGC subtypes, and that RGCs have a different G protein landscape compared to cone photoreceptors. Finally, we demonstrate how microbial opsins, such as GtACR1(*39*), can match and even surpass the light sensitivity of ectopically expressed Opn1sw, suggesting that animal opsins need further study and optimization before becoming reliable optogenetic tools for vision restoration.

Contrary to microbial opsins, which directly exert their effect on membrane potential, for activation vertebrate opsins take advantage of a molecular cascade which includes a heterotrimeric G protein and, at least, a final target channel(*40*). Due to the promiscuous nature of G proteins, previous studies mainly focused on the opsin itself rather than on deciphering the entirety of the cascade being initiated by its activation, the main idea being that since G proteins are present in the targeted cell, their redundant function would lead to neuronal activation upon light stimulation. After observing Opn1sw-mediated hyperpolarization of RGCs, however, it became clear that different ectopically expressed vertebrate opsins can initiate completely different G protein pathways. Since human cells express 4 classes of Ga subunits (Gɑ_s_ (excitatory), Gɑ_i/o_ (inhibitory), Gɑ_q/11_ (excitatory) and Gɑ_12/13_ (excitatory))(*20*), we can speculate that their selective activation could result in opposite effects on cellular polarity in conjunction with different target channels. Therefore, considering GPCR’s selectivity (that is, the preferred sub-class of G proteins preferentially activated by a specific GPCR)(*19*), more attention should be put upfront into the expected effect on cellular polarization when targeting RGCs with vOpsins. While surprising when compared to already published data, the hyperpolarization of RGCs induced by Opn1sw is a closer match to the physiological mechanism taking place in photoreceptors than the MW-opsin-induced depolarization: indeed, rod and cone opsins hyperpolarize the cell membrane upon light stimulation via the inhibitory G protein transducin (belonging to the G_i/o_ family), making the MW-opsin-induced RGC depolarization the more unexpected result. Therefore, a better understanding of the relationship between opsins, G-proteins, intermediate effectors and target channels in ectopic conditions is needed. A step in this direction is represented by the creation of algorithms trained on experimental data and capable of predicting the likeliness of GPCR-G protein interaction solely based on their aminoacidic sequences(*41*). This would, not only streamline the screening process by reducing the amount of experimental testing required, but also allow for a more efficient rational design of novel opsins. In optogenetics for vision restoration, this type of rational design can be found in vertebrate opsin chimeras such as Opto-mGlur6(*38*). Opto-mGlur6 combines the extracellular segment of melanopsin and the intracellular segments of mGlur6, the metabotropic receptor for glutamate found on the membrane of ON-bipolar cells. This design effectively exploits the target cell’s environment to its advantage: instead of having to interact with a novel environment, the opsin (or, better, the intracellular part of the opsin, which is responsible for interacting with the G protein) is already maximally optimized for its task, which in the case of Opto-mGlur6 is interacting with the TRPM1 channel. Going forward, it is clear that approaches based on chimeric opsins(*38*) and alternative bi-stable animal opsins(*42, 43*) hold potential for efficient cellular activation or inhibition. Indeed, a novel opsin from the jellyfish family, named JellyOp, has recently showed good potential for vision restoration(*43*). JellyOp is a GPCR that constitutively binds to Gɑ_s_, enabling extremely fast GIRK responses (34 ± 1 ms) *in vitro* as well as RGC activation *ex vivo* when targeted to ON-BCs. JellyOp also showed a bistable behavior as it could be inactivated by violet light, although at high light intensities (50% GIRK current inactivation at 10^17^photons*s^−1^*cm^−2^).

As mentioned, one of the key reasons for choosing animal opsins over microbial opsins is their higher light sensitivity compared to the one-component microbial opsins. Indeed, early pioneering studies used first and second generation microbial channelrhodopsins(*6, 7, 31, 42–46*) but newly discovered or engineered opsins such as Jaws, GtACR1/2, ChRmine and Chreef(*37, 47, B*), significantly lowered the light intensity threshold required for generating a cellular response closing the gap with ectopically expressed vertebrate opsins. As we show, GtACR1’s sensitivity matches and even surpasses Opn1sw’s in RGCs, showing greater hyperpolarizing responses at intensities as low as ~10^13^ photons*s^−1^*cm^−2^.Several factors could be contributing to such seemingly counterintuitive result. First, while it is true that good levels of expression of membrane-bound Opn1sw can be reached through the SNCG promoter, the density of visual pigment molecules expressed by each cell will inevitably low when compared to the one in photoreceptors’ outer segments(*51*). Furthermore, the different G protein landscape between photoreceptors and inner retinal neurons, resulting in the activation of an alternative and non-optimized G protein cascade with fewer signal amplifying elements between the G protein and the target channel, could result in a lower efficacy. The cascades recruited in different RGC subtypes can also add more variability. Overall, wild-type vertebrate opsins, when ectopically expressed in inner retinal neurons, face a sub-optimal working environment that prevents them from exhibiting their full potential.

Another important aspect when it comes to vision restoration is the opsin’s kinetics, that is, how rapidly it can exert an effect on the membrane potential of a cell. Looking at both our results and the available literature, it is clear that microbial opsins still have the edge over vertebrate opsins. While SNCG-Opn1sw and SNCG-Opn1mw are characterized by a time to peak of about 1100ms and 600ms, respectively (355ms for hSyn-MW-opsin in(*22*)), microbial opsins generally reach their maximum response well below 100 ms after the stimulus onset, no matter the cell type they are expressed in. The faster kinetics of microbial opsins is therefore still more suitable, today, for high refresh, movie-rate vision restoration. In such challenging landscape, however, vertebrate opsins still hold great potential for vision restoration therapies. Going back to chimeric opsins, Opto-mGluR6 achieved highly sensitive, sub-100ms responses at ganglion cell level which almost perfectly matched the physiological, photoreceptor-mediated ones(*38*). The recent optimization of such melanopsin-based chimeras further enhanced vision restoration in mice(*52*).

Overall, in this work we show that despite the current undeniable limitations, vertebrate opsin-based gene therapy still holds promise. We should recognize that the research field is still young, and we are only beginning to scratch the surface of what is possible with these tools. While it is true that unmodified, wild-type human and mouse opsins might not satisfy our needs in terms of kinetics and sensitivity, engineering of novel molecules could bridge the gap with, and maybe surpass, the performance of microbial opsins. Developing a deeper understanding of retinal G protein-related pathways will allow us to accelerate the screening for novel therapeutic molecules, while enhancing current promoters and trafficking of ectopically expressed opsins could result in more efficient outcomes. Finally, insightful molecular engineering of intracellular loops and the opsin’s C-terminal may allow for the creation of faster and more sensitive opsins. Following these paths, vertebrate opsin-based gene therapy may find utility in vision restoration research.

## Acknowledgments

We thank the Paris Vision Institute core facilities (in particular the animal care staff; zoo technicians and regulatory compliance staff of the Animal facility, Histology platform, Imaging platform, and the vector core facility for producing the AAVs; the lab managers; and researchers). We thank financial support from the Foundation Fighting Blindness USA (PPA Grant, Next Generation Optogenetics to DDA), Deutsche Forschungsgemeinschaft (DFG) grants: He2471/23-1 (to S.H.), He2471/21-1 (to S.H.), Project ID 316803389—SFB 1280 project A07 (to S.H.), Project ID 492434978—GRK 2862/1 projects 01 (to S.H.). We acknowledge LabEx LIFESENSES (ANR-10-LABX-65) and IHU FOReSIGHT (ANR-18-IAHU-01), DIM C-BRAINS, funded by the Conseil Régional d’Ile-de-France, Région Ile-de-France and Paris Ile-de-France Region under « DIM Thérapie génique » initiative.

