## Supplemental Figure 1 for "Ectopic expression of two cone opsins in mouse RGCs results in opposite responses to light stimulation, likely due to differential G protein activation"

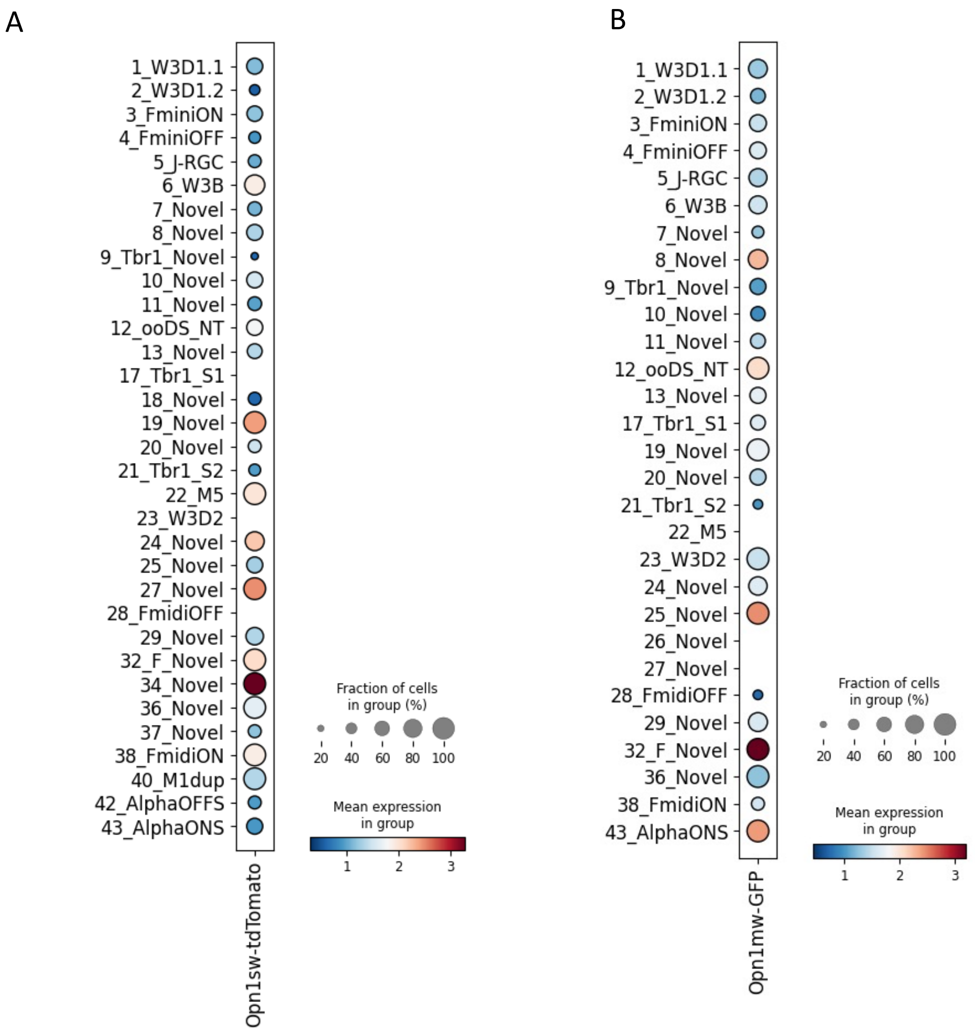


**Supplementary Figure 1**. **Transgene expression across RGCs population** A) in SNCG-Opn1sw B) in SNCG-Opn1mw-injected (right) rd1 retinas. In both conditions, the SNCG promoter does not seem to discriminate between RGC subtypes, suggesting that the variability in cellular response does not originate from a biased transgene expression in a particular RGC subtype.

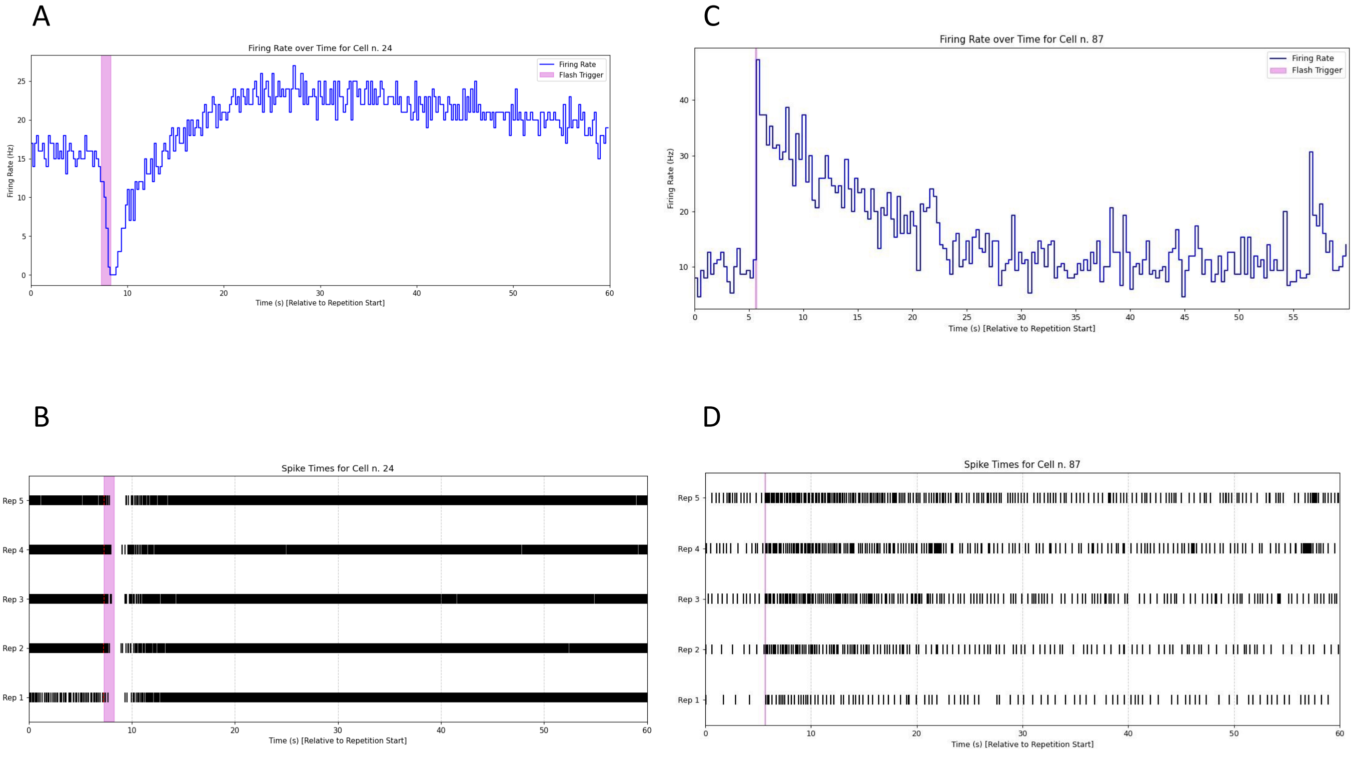


**Supplementary Figure 2. Spike sorting on a SNCG-Opn1sw-injected retina.**

**A**) Representative trace of a strongly hyperpolarizing RGC. **B) R**aster plot showing the response of the RGC to 5 flashes of 1s at 380nm light and 3.5e+14 photons*cm$^{2}$*s$^{-1}$. **C)** Representative trace of a depolarizing RGC; **D)** Raster plot showing the response of the RGC to 5 flashes of 100ms at 535nm light and 2.3e+14 photons*cm$^{2}$*s$^{-1}$. Bin size for the psth was set to 200ms in both A and C.

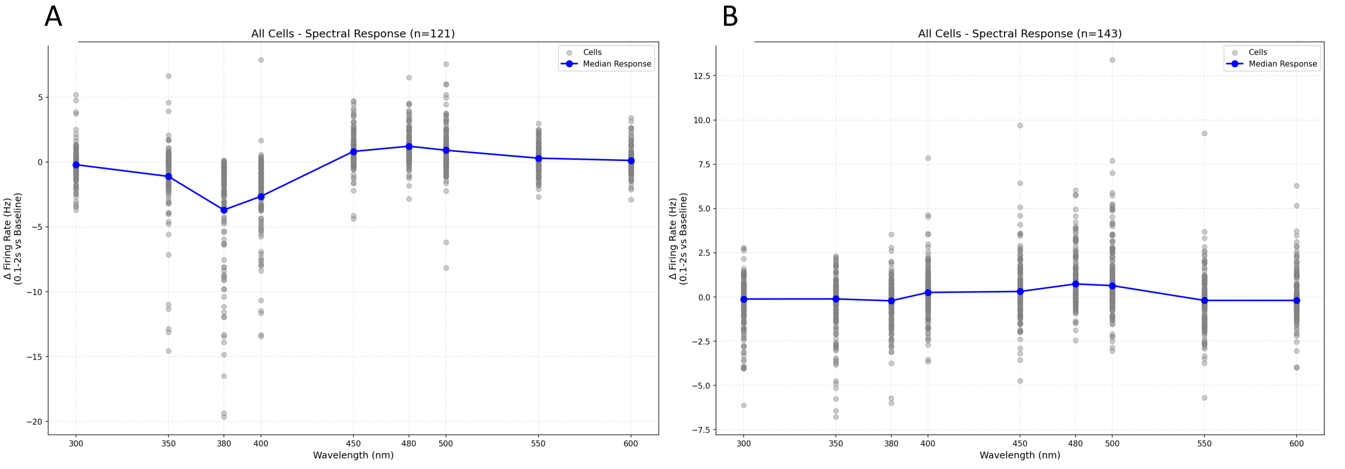


**Supplementary Figure 3. Spectral analysis of sorted cells**. (A-B) Spectral analysis conducted on single cells from (A) SNCG-Opn1sw-injected rd1 retinas confirms that hyperpolarizing responses are primarily generated at 380nm, with detectable responses at 400nm; (B) hyperpolarizing responses are not observed in non-injected rd1 retinas. Cell responses were averaged over 5 flashes of 1s for each wavelength. Light intensity: approx. 3.5e+14 photons*$cm^2$*$s^{-1}$.
